# Representation of motion concepts in occipitotemporal cortex: fMRI activation, decoding and connectivity analyses

**DOI:** 10.1101/2021.09.30.462531

**Authors:** Yueyang Zhang, Rafael Lemarchand, Aliff Asyraff, Paul Hoffman

## Abstract

Embodied theories of semantic cognition predict that brain regions involved in motion perception are engaged when people comprehend motion concepts expressed in language. Left lateral occipitotemporal cortex (LOTC) is implicated in both motion perception and motion concept processing but prior studies have produced mixed findings on which parts of this region are engaged by motion language. We scanned participants performing semantic judgements about sentences describing motion events and static events. We performed univariate analyses, multivariate pattern analyses (MVPA) and psychophysiological interaction (PPI) analyses to investigate the effect of motion on activity and connectivity in different parts of LOTC. In multivariate analyses that decoded whether a sentence described motion or not, the middle and posterior parts of LOTC showed above-chance level performance, with performance exceeding that of other brain regions. Univariate ROI analyses found the middle part of LOTC was more active for motion events than static ones. Finally, PPI analyses found that when processing motion events, the middle and posterior parts of LOTC (overlapping with motion perception regions), increased their connectivity with cognitive control regions. Taken together, these results indicate that the more posterior parts of LOTC, including motion perception cortex, respond differently to motion vs. static events. These findings are consistent with embodiment accounts of semantic processing, and suggest that understanding verbal descriptions of motion engages areas of the occipitotemporal cortex involved in perceiving motion.

## Introduction

Embodied theories of semantics hold that we represent knowledge of concepts by simulating the sensory, motor and other sensations they elicit (Barsalou, 1999; Decety & Grezes, 2006; Gallese & Lakoff, 2005). In terms of motion concept representation, there is evidence that brain areas involved in perceiving and controlling movements are also engaged when we process concepts relating to motion (Barsalou, 2003; Beilock et al., 2008). In particular, the lateral occipital-temporal cortex (LOTC) has been implicated in processing, perceiving and representing embodied experiences of perceived motion (Lingnau & Downing, 2015; Tucciarelli et al., 2019). The LOTC is typically assumed to encompass the posterior portion of the middle temporal gyrus, extending back into middle occipital gyrus as far as the lateral occipital sulcus (Lingnau & Downing, 2015; Weiner & Grill-Spector, 2013) (see Figure 1). This broad region of the cortex includes areas implicated in motion and action perception as well as sites associated with language processing. As an important area for action perception, parts of LOTC respond when participants watch videos or pictures of human body movements (Cross et al., 2006; Lingnau & Petris, 2013), tool-related actions (Beauchamp et al., 2002), and abstract moving stimuli formed by dots (Tootell et al., 1995; Wall et al., 2008) or geometries (Zeki et al., 1991). In addition, lesion studies have showed that damage to LOTC leads to poor performance in naming, preparing or imitating actions (Brambati et al., 2006; Buxbaum et al., 2014; Hoeren et al., 2014). Body representation is also a function of LOTC, with posterior parts of the region identified as ‘limb-selective’ regions (Weiner & Grill-Spector, 2013).

**Figure 1.**
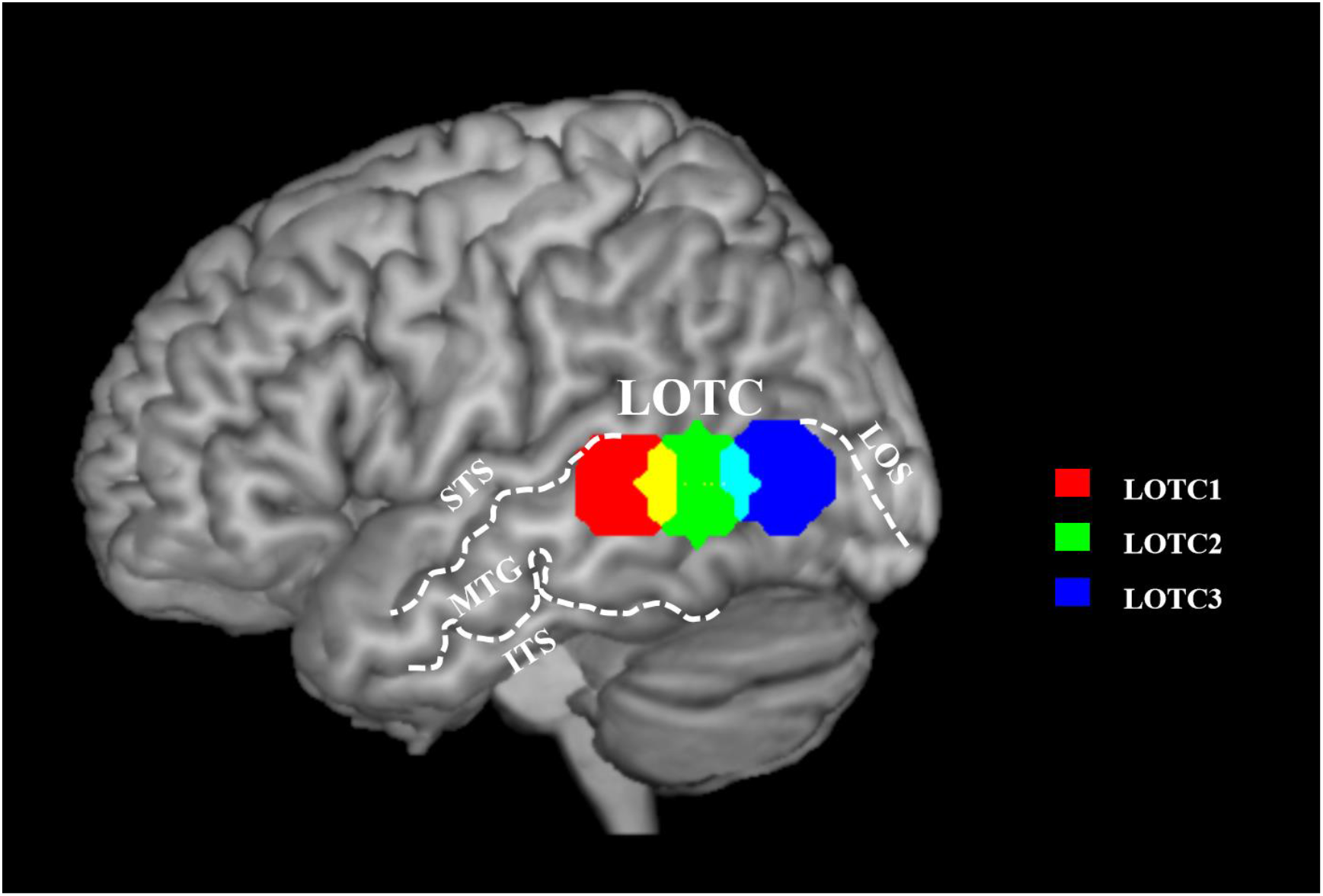
The lateral occipital temporal cortex and its division into regions of interest for this study. ITS, inferior temporal sulcus; MTG, middle temporal gyrus; STS, superior temporal sulcus; LOS, lateral occipital sulcus.

Beyond perception, understanding action/motion concepts also engages LOTC. Some positron emission tomography (PET) studies and functional magnetic resonance imaging (fMRI) studies have observed LOTC activity when participants generate appropriate verbs for objects (Martin et al., 1995) or match motion pictures with similar semantic meanings (Tucciarelli et al., 2019). Lesion studies have also reported that patients with damage in LOTC showed deficits of matching verbal descriptions of actions to related pictures (Kalénine et al., 2010; Kemmerer et al., 2012; Urgesi et al., 2014). Some researchers suggested that the response of LOTC is also sensitive to grammatical categories. For example, compared with nouns, verbs elicit stronger activation in parts of LOTC (Bedny et al., 2014; Kable et al., 2005). Within verbs, Peelen et al. (2012) found that the posterior part of LOTC showed a preference for action verbs (verbs describing dynamic activities, like ‘to walk’) than state verbs (verbs describing state or mental actions, but no obvious body actions, like ‘to believe’).

These results have shown that LOTC includes areas involved in both perceiving action and motion visually and in understanding related semantic concepts. However, there remains considerable debate over the degree to which the precise regions engaged by these functions overlap and consequently, researchers hold different views about the role of LOTC in motion concept representation. Strong re-enactment theories claim that understanding motion words requires reactivation of corresponding perceptual experiences, and thus predict that understanding motion words elicits similar neural responses to perceiving motion directly (Hauk et al., 2004; Kiefer et al., 2012; Pulvermüller, 2005; Saygin et al., 2010). Weaker embodiment theories argue that representation of motion concepts recruit regions close to relevant sensory areas, but do not necessarily activate the perceptual regions themselves (Barsalou, 2003; Bedny et al., 2008; Kable et al., 2002; Martin & Chao, 2001). Finally, modality-independent views propose that LOTC activation for motion words is driven by retrieval of event concepts or grammatical information linked with verbs, rather than effects of sensory-motor simulation (Bedny & Caramazza, 2011; Bedny et al., 2012).

To understand the role of LOTC in action/motion concept representation, it is critical to determine whether processing motion-related words engages the same parts of LOTC, in the same ways, as perceiving motion directly. Evidence on this issue has been somewhat inconsistent. Many studies have focused on area V5 (also frequently termed hMT+), a critical region for perception of visual motion located in the posterior part of LOTC. As an important region for encoding visual information concerning motions and body movements (Beauchamp et al., 2004; Dumoulin et al., 2000; Kourtzi & Kanwisher, 2000; Liu et al., 2016; Schultz et al., 2005; Thompson et al., 2005), V5 was also found to activate during action concept processing (Assmus et al., 2007; Glenberg & Kaschak, 2002; Revill et al., 2008; Rueschemeyer et al., 2010; Saygin et al., 2010). It was reported that, compared with processing language or images of static events, more activation of V5 was observed when participants read and listened to sentences describing motion events (Rueschemeyer et al., 2010; Saygin et al., 2010), made semantic decisions to words (Kable et al., 2002) or sentences (Revill et al., 2008) describing motion, or comprehended action knowledge represented in static pictograms (Assmus et al., 2007).

However, studies finding V5 activation have been criticized for using pictures as stimuli (Assmus et al., 2007; Kable et al., 2002), or for combining motion-related language stimuli with other visual stimuli (Revill et al., 2008; Rueschemeyer et al., 2010; Saygin et al., 2010). Since pictures of static objects or humans can also elicit responses in V5 (Kourtzi & Kanwisher, 2000; Senior et al., 2000), it is hard to determine the contribution of conceptual processing when language and pictures are presented together. In addition, integrating audio-visual language stimuli could activate V5 (Calvert et al., 1999; 2000), thus the V5 effect in some studies might be caused by the use of audio-visual stimuli (Saygin et al., 2010), instead of the motion content.

In addition, not all studies have supported the view that V5 is engaged when processing motion-related words. Some evidence suggests that only the more anterior part of LOTC, typically referred to as pMTG (posterior middle temporal gyrus), shows such effects. Several studies using pure language stimuli have only found motion effects in areas anterior to V5, such as pMTG (Bedny et al., 2008; Gennari et al., 2007; Noppeney et al., 2005). For instance, Bedny et al. (2008) asked participants to judge the semantic similarity of words (nouns and verbs), finding more activation for verbs in pMTG but not V5. Kable et al. (2002) used a conceptual matching task (matching related words or pictures) and observed that both pMTG and V5 responded to motion images, but only pMTG and other semantic processing regions were activated for motion words. These indicated that representing motion concepts might not recruit V5 directly, a view supported by a meta-analysis by Watson et al. (2013), which reported that activation for action verbs was more anterior in LOTC than for action images.

The absence of V5 effects has also been reported in studies using motion and static sentences as stimuli (Chen et al., 2008; Desai et al., 2013; Dravida et al., 2013; Humphreys et al., 2013; Wallentin et al., 2005). When comparing motion sentences (e.g ‘The child fell under the slide’) with static or abstract ones (e.g ‘The merchant was greedy’ / ‘The congress is causing a big trade deficit again’), these studies found stronger responses in the pMTG region anterior to V5, but not in V5 itself. In particular, Humphreys et al. (2013) separated motion sentences from static images depicting motion in every trial. They showed participants a picture depicting a moving or static event first, followed by a recording of a sentence describing the event presented after a short interval. They reported that V5 responded to the motion pictures, but not to the later sentence stimuli. These findings have made some researchers skeptical about the role of lower-level perceptual areas such as V5 in understanding motion language (Gennari, 2012; Humphreys et al., 2013).

It is important to note that the studies reviewed above relied on univariate neuroimaging analyses, in which data from each voxel is analyzed independently. Multi-voxel pattern analysis (MVPA) has the potential to provide more sensitive analyses of motion effects in LOTC and to uncover the content of representations in this brain region, but has only recently been applied to this research question. Wurm and Caramazza (2019) used MVPA to investigate representation of different types of motion elicited by videos and sentences (e.g., “the girl opens the bottle”). They found that regions in LOTC encoded motion types in a crossmodal fashion (generalizing between videos and sentences), supporting the general view that this area is involved in conceptual representation of motion and action. However, these authors did not directly compare effects in posterior LOTC (V5) with those more anterior parts (pMTG), thus the spatial distribution of these effects remains unclear. Other MVPA studies using video stimuli have revealed that both concrete and abstract action representations can be decoded from fMRI signal in posterior LOTC (Wurm et al., 2016), and that the neural responses to observed actions can be classified in terms of transitivity and sociality (Wurm et al., 2017). However, no studies have yet investigated how MVPA effects to pure linguistic descriptions of motion vary across the LOTC region.

In summary, while it is clear that LOTC is engaged by motion understanding as well as motion perception, the precise locus of these effects, and hence their interpretation, remains under debate. In present study, we used multiple analysis methods to construct a more detailed picture of how the response to motion sentences varies across LOTC. We re-analysed data collected by Asyraff et al. (2021), in which participants were asked to make simple semantic decisions of sentences describing 4 events. We categorized the 4 events as 2 motion events and 2 static events. We used univariate analysis, MVPA, and psychophysiological interaction (PPI) analysis to explore activation patterns within LOTC and its functional connectivity with other areas. We used sentences rather than words or phrases as stimuli, as sentences are more likely to elicit strong sensory-motor imagery (Dravida et al., 2013). We planned to ask three questions. The first is similar to those in previous univariate studies: Which areas within LOTC activate more to motion sentences than to static ones? Going beyond this, however, we used MVPA to investigate the degree to which neural patterns in LOTC could distinguish motion from static sentences as well as between different forms of motion. This allowed us to ask what level of motion knowledge is represented in LOTC during sentence processing. Finally, we used PPI to investigate how connectivity between LOTC and other brain regions changed as a function of sentence type. This allowed us to gain additional information about the role of LOTC regions by revealing their interactions with other neural systems. To guide our analyses, we divided LOTC into three sub-regions along its anterior to posterior axis (see Figure 1), guided by peak meta-analytic activations for semantics and motion processing. By doing this, we aimed to establish a better understanding of how function varies across the LOTC region during processing of motion language.

## Method

### Participants

26 healthy participants were recruited (20 female; mean age 22.48, range 18-35 years). All participants were right-handed native English speakers, and no one reported history of dyslexia or other neurological disorders. The study was approved by University of Edinburgh School of Philosophy, Psychology & Language Sciences Research Ethics Committee. Other analyses of the data presented here have been reported by Asyraff et al. (2021). This previous study used MVPA to investigate semantic representation of general event concepts; here, we specifically investigated differences between motion and static sentences, with a focus on LOTC.

### Materials

32 sentences were created as stimuli. Half of these (16) were target stimuli and the other half were fillers. The target sentences described four different events, each with a different agent, patient and verb (see Table 1). Two events involved an agent making a visualizable movement in relation to the object, while the other two involved static acts. Four descriptions of each event by substituting lexical terms with similar meanings (e.g. ‘cow’ and ‘bull’) and by varying the syntactic structure of the sentence (active vs. passive). The 16 fillers were anomalous sentences created with words used in target stimuli, and with the same syntactic structures. Fillers did not describe a coherent, meaningful event (e.g. The computer jumped over the bull).

**Table 1:** Target stimuli used in the experiment.

| Condition | Event | Syntactic form | Lexical items 1 | Lexical items 2 |
| --- | --- | --- | --- | --- |
| Motion | Event1 | Active | The bull leapt over the gate. | The cow jumped over the fence. |
|  |  | Passive | The gate was leapt over by the bull. | The fence was jumped over by the cow. |
|  | Event2 | Active | The lorry bumped the lamp post. | The truck hit the street light. |
|  |  | Passive | The lamp post was bumped by the lorry. | The street light was hit by the truck. |
| Static | Event3 | Active | The computer processed the file. | The laptop analysed the document. |
|  |  | Passive | The file was processed by the computer. | The document was analysed by the laptop. |
|  | Event4 | Active | The student considered the problem. | The pupil pondered the issue. |
|  |  | Passive | The problem was considered by the student. | The issue was pondered by the pupil. |

In Asyraff et al. (2021), all 32 sentences were rated by 18 participants who did not take part in the main experiment. A five-point scale was used for rating how meaningful a sentence was; target stimuli received significantly higher scores than fillers (Target M = 4.56, SD = 0.32; Filler M = 1.53, SD = 0.57; t(30) = 18.6, p<0.001). For the current study, in order to verify our assignment of sentences as motion events and static events, the 16 target sentences were rated with a seven-point scale by 22 native English speakers (who did not participate in the main experiment) for the degree to which each sentence brings to mind an experience of motion. Motion events received significantly higher scores than static events (Motion M = 5.42, Static M = 2.23, mean difference = 3.18, t(9.66)=17.76, p<0.0001). Raters also rated each sentence for visual, auditory and emotional experiences. No difference for found for emotion (mean difference = 0.24, t(12.23)=0.67, p=0.52); however, motion events were more associated with visual (mean difference = 2.23, t(13.23)=11.87, p<0.0001) and auditory experiences (mean difference = 1.92, t(7.99)=7.84, p<0.0001). Importantly, the difference between conditions was considerably larger for the motion ratings than for either visual or auditory. Thus, the perception of motion was the most salient difference between our sets of stimuli, but motion events also brought to mind richer auditory and visual experiences for participants, in line with the real-word perceptual correlates of motion.

We also computed word frequency and concreteness for every target sentence by averaging values of the content words in each sentence. Concreteness values were obtained from Brysbaert et al. (2014) and frequency values from Van Heuven et al. (2014). There was no significant difference in word frequency (Motion M=5.69, Static M=5.72, t(11.77)=-0.025, p=0.89) but motion events were described with more concrete words than static events (Motion M =3.04, Static M =2.54, t(13.27)=3.65, p<0.01). This reflects the fact that motion verbs are easier to visualize than static verbs. Importantly, neuroimaging meta-analyses indicate that activation differences between concrete and abstract concepts are not typically observed in LOTC (Bucur & Papagno, 2021; Wang et al., 2010). Thus effects observed in this study are unlikely to be due to the concreteness difference between motion and static events. The list of all sentences and their properties can be found in Supplementary Materials.

### Experiment procedure

On each trial, a fixation cross was presented for 500ms, followed by one of the 32 sentences presented in the centre of the screen for 4000ms. Participants were required to judge whether the sentence was meaningful by pressing buttons held in the left and right hands. The order of sentence presentation was randomized separately for each participant in each run. By fully randomizing presentation orders for each run and participant, we ensured that there was no temporal structure present in the data that could lead to false positive errors during MVPA classification (Mumford et al., 2014). After each trial, there was a jittered interval of 4000ms to 8000ms. The length of the interval was randomized independently of sentence order randomization. Each run presented all 32 sentences once and each participant was asked to complete 6 runs in total. We note that, because the same sentences were used in each run, neural responses are potentially subject to the repetition suppression effect, whereby activation decreases when the same stimuli are processed repeatedly (Barron et al., 2016). There are two reasons why we do not believe this poses a problem for the present study. First, repetition suppression effects in fMRI are short-lived, dissipating on the order of seconds, and are strongest when few other stimuli are presented between repetitions (Barron et al., 2016). In the present study, stimulus repetitions were separated by a mean of 32 stimuli and 360s (the length of one run). Second, motion and static sentences were repeated equally often, so any repetition suppression should affect both sentence types equally.

For the behavioral data, T-tests were used to compare reaction time and accuracy for motion vs. static sentences. In addition, R-4.0.3, with the ‘lme’, ‘effects’ and ‘afex’ packages, was employed to build a linear mixed effect model predicting RTs on trials with correct responses. Fixed effects included the type of events (motion/static), run number in the experiment and sentence length (number of characters in each sentence). Sentence length was set as a fixed effect as sentences were not precisely matched for length. Participant was set as the random effect with intercepts and random slopes for event types.

### Data acquisition

A 3T Siemens Prisma scanner and 32-channel head coil were employed for the data acquisition. The T1-weighted structure scan was acquired with TR = 2.62s, TE = 4.5ms and 0.8mm^3^ voxels. For functional scanning, each image contained 46 slices of 3mm^3^ isotropic voxels with an 80*80 matrix and the TR for scanning was 1.7s. To improve fMRI signal quality and reduce the influence caused by movement and other artefacts, a whole-brain multi-echo acquisition protocol was used. The signal was collected at 3 echo times (13ms, 31ms, 48ms) simultaneously (Feinberg et al., 2010; Moeller et al., 2010; Xu et al., 2013), then the images were weighted and combined and independent components analysis (ICA) was used to remove noise components.

### Preprocessing

SPM12 and TE-Dependent Analysis Toolbox 0.0.7 (Tedana) (DuPre et al., 2019) were employed for preprocessing. Slice-timing correction and motion correction were performed first. The data acquired at the first echo time (13ms) was used for estimating motion parameters (Power et al., 2017). Next, we used Tedana to combine the data acquired at the 3 echo times into one single time-series and denoise images in each run (Kundu et al., 2017). Tedana uses ICA to discriminate noise and task-related signal, based on their different patterns of signal decay over increasing TEs. Finally, we employed SPM12 to coregister functional scans to the anatomical images and normalise them to MNI space with DARTEL (Ashburner, 2007).

For univariate and PPI analyses, images were smoothed with a kernel of 8mm FWHM. We processed data with a high-pass filter with a cut-off of 128s and used one general linear model for analyzing all 6 runs. In a model of a run, 3 regressors modelled motion events, static events and anomalous events respectively. Covariates consisted of six motion parameters and their first-order derivatives.

### Regions of Interest

We divided left LOTC into 3 sub-regions horizontally: LOTC1, LOTC2 and LOTC3 (Figure 1). Each sub-region was defined as a 5mm radius sphere centered on specific MNI co-ordinates. Co-ordinates for LOTC1 and LOTC3 were obtained from automated meta-analysis of the neuroimaging literature using the Neurosynth database (Yarkoni et al., 2011).

The centre of LOTC1 was located at [-52 -40 2], the peak co-ordinate in LOTC for activations associated with the term “semantic” in Neurosynth (see Figure 2A). Thus, LOTC1 represented an anterior location within LOTC associated with general semantic processing. Studies of semantic processing frequently refer to this area as pMTG. In contrast, LOTC3 was centered at [-44 -72 5], the peak co-ordinate from activations associated with the term “motion” in Neurosynth (see Figure 2A). Figure 2B shows the overlap between LOTC3 and area V5, as defined in the probabilistic atlas of Malikovic et al. (2007). In addition, the centre co-ordinates of LOTC3 are very close to those obtained in studies that used motion perception tasks to localize this area (Dravida et al., 2013; Humphreys et al., 2013; Saygin et al., 2010). Finally, for LOTC2 we needed to identify a region located between the anterior and posterior extremes of LOTC1 and LOTC3. We selected the peak co-ordinate from the whole-brain analysis of meaningful sentences > rest in our own study [-54 -55 3], as this fell midway between the other two regions. Note that the use of this peak did not bias us towards finding any particular effect of motion over static sentences, since the meaningful sentences condition contained equal numbers of both types of sentence. The 3 sub-regions overlapped each other slightly so that we could plot graded changes in the functional profile of LOTC along its anterior-to-posterior axis. Together they covered the entire cortical territory thought to comprise LOTC.

**Figure 2.**
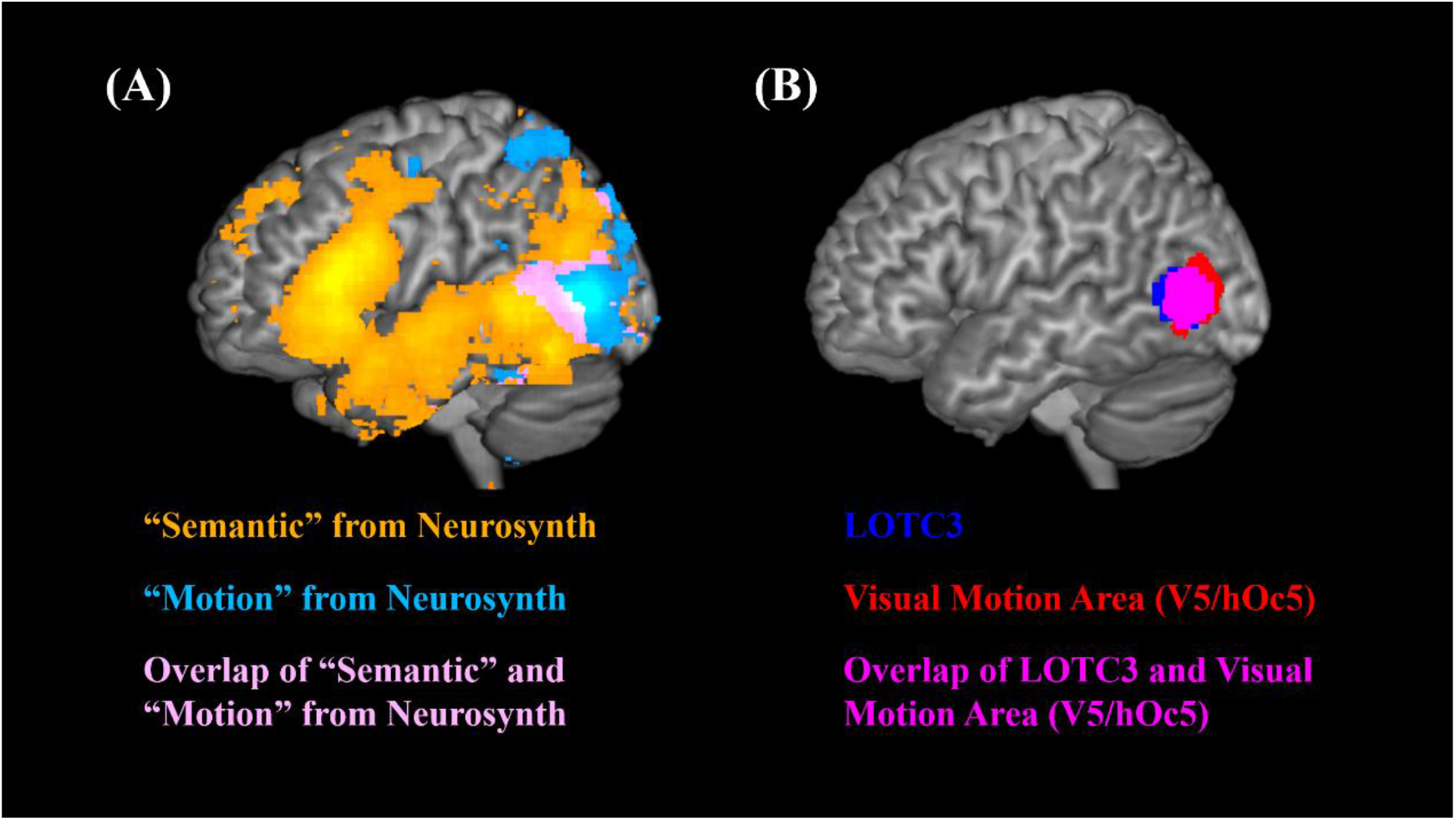
Details about ROI definitions. (A). Regions significantly associated with “Semantic” and “Motion” keywords in Neurosynth, (FDR p < 0.01); (B). Overlap of LOTC3 and Area V5. V5 areas are those exceeding 40% probability of falling within V5/hOc5 in a probabilistic map (Malikovic et al., 2007)

### Univariate fMRI analyses

Both whole-brain analysis and ROI analyses were conducted, contrasting motion and static sentences. The whole-brain analysis was corrected for multiple comparisons (*p* < 0.05) at the cluster level using SPM’s random field theory, with a cluster-forming threshold of *p* < 0.005. For the univariate ROI analyses, we first used SPM12 to extract the mean beta values of LOTC1, LOTC2 and LOTC3 in motion event and static event conditions, which represent activation relative to the implicit baseline (rest). The beta values were imported into R-4.0.3, and an ANOVA analysis was completed to assess effects of event type (motion, static), ROI and their interaction.

### Multivariate pattern analysis (MVPA)

We employed MVPA to investigate which areas discriminate between different types of event. Since some studies have reported that smoothing slightly improves performance in decoding models (Gardumi et al., 2016; Hendriks et al., 2017), images used for MVPA were normalized and smoothed at 4mm FWHM. To obtain T-maps for each of the 32 sentences, a general linear model was built for each run. In each model, each sentence was modelled with a separate regressor. We used CoSMoMVPA (Oosterhof et al., 2016) for processing the T-maps generated by these models.

Previous studies have shown that MVPA can be very sensitive to differences in reaction time between classes (Todd et al., 2013; Woolgar et al., 2014). In the present study, participants were reliably slower to respond to motion sentences when compared with static sentences (see Results). To avoid the possibility that this difference could lead to successful classification, we regressed out the effect of RT on each voxel’s t-values prior to MVPA. We did this by estimating a linear model for each voxel predicting t-values from RT on a trial-by-trial basis. The residuals of these models, which were uncorrelated with RT, were used as the patterns in the classifier (Todd et al., 2013).

Three analyses were performed, repeated at the searchlight and ROI level (with the ROIs defined earlier). For the first analysis, a decoding model was trained to classify whether activation patterns belonged to motion or static sentences. This analysis was intended to reveal which regions were sensitive to the presence of motion in event descriptions. For the other 2 analyses, we trained 2 models separately to discriminate between the 2 kinds of motion events and the 2 kinds of static events used in the study. These 2 analyses were intended to identify areas that coded for the semantic content within the motion and static domains. Details of each decoding model are as follows:

### Decoding event types (motion vs. static)

An example of train and test patterns for this classifier is shown in Figure 5B. Because each event could be described using four different sentences (differing in lexical or syntactic forms), we were able to test the ability of activation patterns to classify motion in a way that generalizes to novel sentences. For one iteration, the classifier was trained on 3 of the 4 sentences describing each event and tested on the remaining sentences (one for each event) that were not used for training. This process was repeated 16 times, until all possible combinations of the various lexical forms and syntactic forms were used as the training set. We adopted this “leave-one-stimulus-out” approach, as opposed to the more common “leave-one-run-out” method, because it provides a stronger test of our hypothesis. Specifically, requiring generalization to novel sentences ensures that successful decoding is driven by conceptual content and not by lower-level characteristics of particular stimuli (Asyraff et al., 2021). The first searchlight analysis gave us a whole-brain decoding accuracy map (Figure 5A). The ROI analysis used patterns in LOTC1, 2 and 3 for decoding (Figure 7A), and tested whether these accuracies were significantly higher than 50% (chance level for classifying two categories).

### Decoding specific motion events

A example of one iteration in this analysis can be seen in Figure 6B. The training sets and testing sets were partitioned in a way similar way to decoding event types, except that only motion sentences were used and the classifier was required to discriminate between the two events that involved motion. The process was also repeated 16 times, until all possible combinations of different event descriptions had been used as the training set.

### Decoding specific static events

This took the same form at decoding specific motion events, except that the classifier was trained to discriminate the two static events. An example of one iteration can be seen in Figure 6D.

All classifiers were trained with a support vector machine (LIBSVM) with the regularization parameter *C* set to 1. To test whether the models could classify better than chance level, we used a two-stage method to perform permutation tests (Stelzer et al., 2013). Specifically, a decoding model was trained and tested 100 times for each participant (divided equally between all iterations of the training set), with the class labels randomly permuted in each run. This process provided a distribution of accuracies under the null hypothesis for each participant. Then we used a Monte Carlo approach to compute a null accuracy distribution at the group level (over all participants). From each participant’s null distribution, we selected one random accuracy value/map for each training iteration and averaged them to generate a group mean. This process was repeated 10,000 times to generate a distribution of the expected group accuracy under the null hypothesis. In searchlight analyses, we entered the observed and null accuracy maps into the Monte Carlo cluster statistics function of CoSMoMVPA to generate a statistical map corrected for multiple comparisons using threshold-free cluster enhancement (Smith & Nichols, 2009). These maps were thresholded at corrected *p* < 0.05. For ROI analyses, we used the position of the observed group accuracy in the null distribution to determine the p-value (e.g. if the observed accuracy was greater than 95% of accuracies in the null distribution, the p-value would be 0.05).

### Psychophysiological interaction (PPI) analyses

PPI analysis is a method for investigating task-specific changes in the relationship between different brain regions’ activity (Friston et al., 1997). PPI is a form of functional connectivity analysis. However, while functional connectivity analyses often consider the temporal correlations between different brain regions in all conditions (including the resting state), PPI focuses specifically on *changes* in connectivity caused by experimental manipulations (Ashburner et al., 2014; Gitelman et al., 2003; O’Reilly et al., 2012). In current study, we used PPI analysis to investigate which brain regions’ activity would show increased correlation with LOTC sub-regions during processing of motion (relative to static) sentences. The PPI analysis for each seed region (LOTC1, LOTC2, LOTC3) was conducted using SPM12 with the following steps. First, the seed region was defined as described in the Region of Interest section above, and the BOLD signal time-series from the seed region was extracted using the first eigenvariate. Then, a general linear model was built with the following regressors:

1. The signal in the seed region.
2. A regressor coding for the experimental effect of interest, where motion sentence trials were coded as ‘+1’ and static sentences ‘-1’.
3. The interaction between the signal in the seed region and the experimental effect (motion/static).
4. An additional regressor coding for the presentation of anomalous sentences, as a nuisance covariate.
5. Head movement covariates as included in the main univariate analysis.

This model was used for testing effects of the PPI regressor (i.e., changes in connectivity driven by sentence type) in the whole brain. Results were corrected for multiple comparisons (*p* < 0.05) at the cluster using SPM’s random field theory, with a cluster-forming threshold of *p* < 0.005.

### Data and code availability

Group-level results maps and ROI masks are archived at: https://neurovault.org/collections/11009/. Other study data and code are available at: https://osf.io/d3nkc/.

## Results

### Behavioural data

Paired t-tests were conducted to examine whether participants responded differently to static vs. motion events. No significant difference was found in their accuracies (static M = 93.35%, SD = 0.09, motion M = 95.11%, SD = 0.08, t(25) = -1.06, p < 0.3), but participants reacted faster when processing static events (static M = 1799ms, SD = 452.02, motion M = 1931ms, SD = 459.91, t(25) = -4.27, p < 0.0002). In a linear mixed effects model controlling for the effect of run order and length of sentences, the event type still had a significant effect on reaction time (t(2267)=-7.316, p< 0.001).

### fMRI data

Three sets of analysis were formed on the fMRI data: univariate analyses, multivariate pattern analyses (MVPA) and psychophysiological analyses (PPI).

### Univariate analyses

In the whole-brain analysis contrasting motion vs. static sentences, there were no significant differences with cluster FWE correction. However, some small clusters of activation for motion > static events were observed at uncorrected p<0.005 (Figure 3). These included a small cluster in LOTC and clusters in regions of parahippocampal gyrus and lateral occipital cortex.

**Figure 3.**
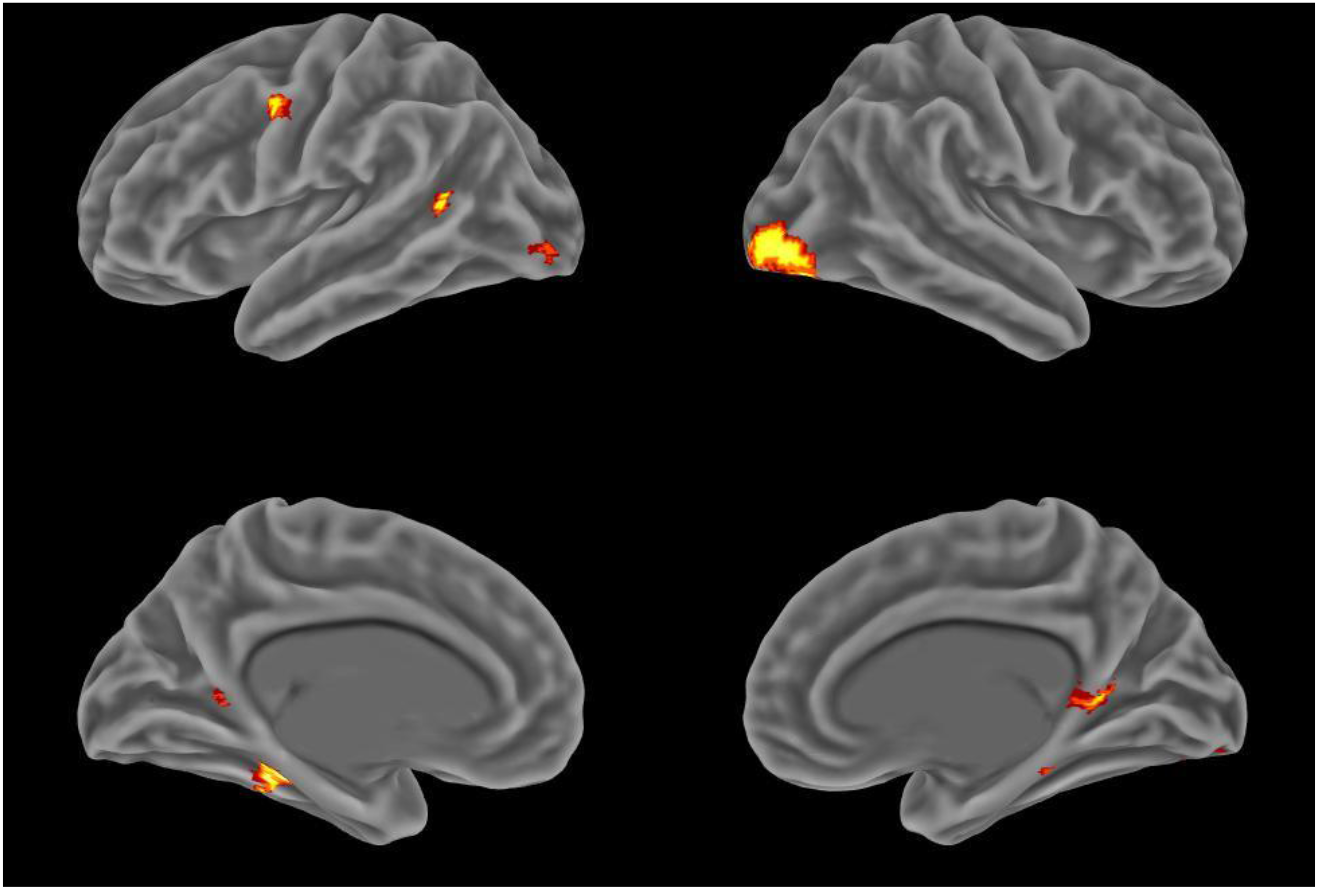
Univariate effects of motion events minus static events at a liberal threshold (p<0.005, uncorrected)

Figure 4 shows activation estimates for the contrast of motion and static events in the three ROIs comprising LOTC. We used two-way repeated measures ANVOA to test whether the effect of motion varied across the sub-regions of LOTC. The analysis revealed a significant main effect of sub-region (F(1.43, 35.83) = 13.057, p<0.001) and an interaction between sub-region and condition (F(2, 50) = 5.947, p<0.005). Post-hoc tests in each ROI found greater activation for motion events in LOTC2 only (t(25)=2.56, p<0.017). A comparison of all meaningful sentences vs. rest revealed activation in LOTC1 and LOTC2 (see Supplementary Figure 1).

**Figure 4.**
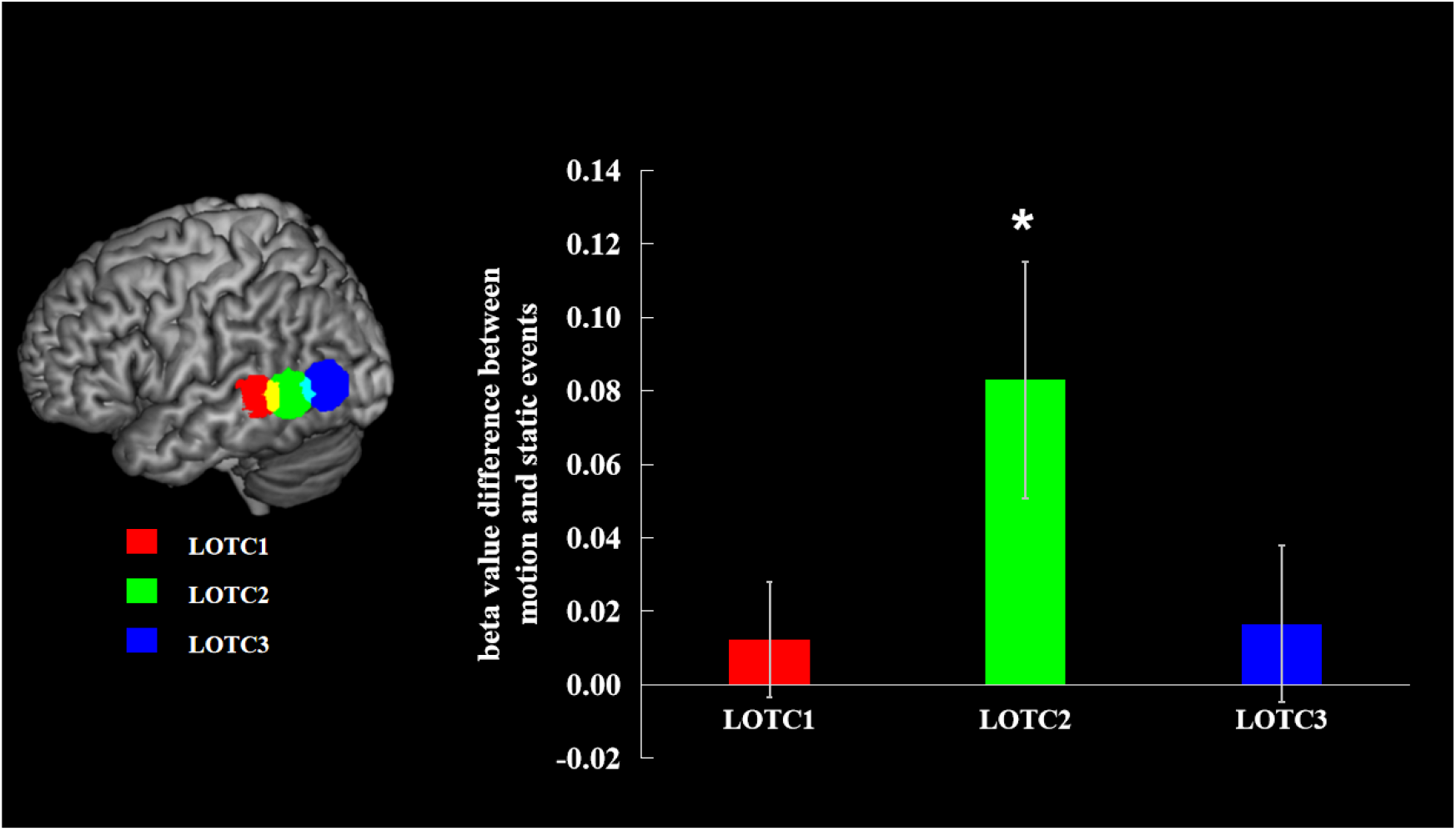
Effects of motion minus static events in LOTC ROIs. Bars show one standard error of the mean.

### MVPA analyses

Figure 5 displays the result of the first analysis, which discriminated between motion and static events. Coloured areas indicate regions where decoding accuracy significantly exceeded chance levels (cluster-corrected p<0.05). Successful decoding was achieved in left hemisphere temporal and occipital regions. The highest decoding accuracy was found in left LOTC. The right lingual gyrus also showed above-chance level decoding accuracy.

**Figure 5.**
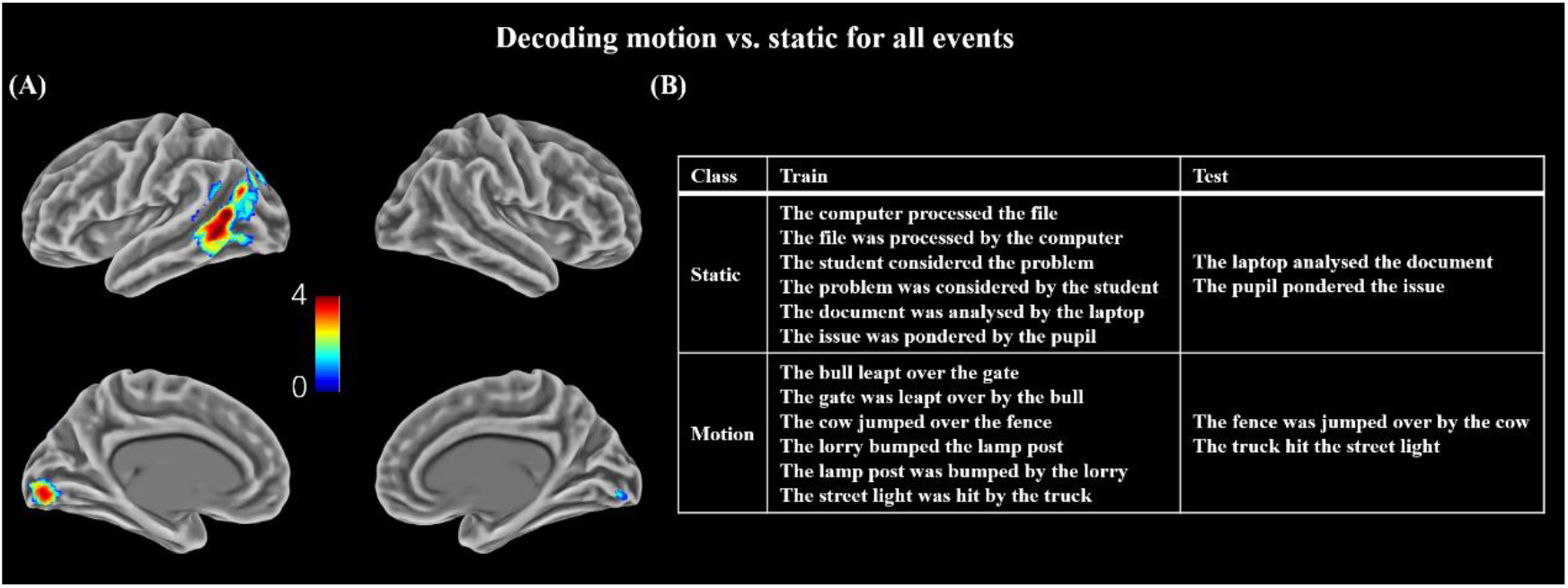
Decoding of event types (motion vs. static). (A) Decoding accuracy map, showing regions where classifier performance exceeded chance (corrected p < 0.05). (B) Example of train and test patterns for one iteration of the analysis.

Figure 6 shows the results of classifiers trained to discriminate between the two motion events and the two static events used in the study. For the static events, successful decoding was observed in a wide range of regions in both hemispheres, including LOTC. However, parts of left occipital and parietal cortex showed the best decoding performance. Discrimination between the two motion events was unsuccessful: above-chance decoding was not observed anywhere in the brain.

**Figure 6.**
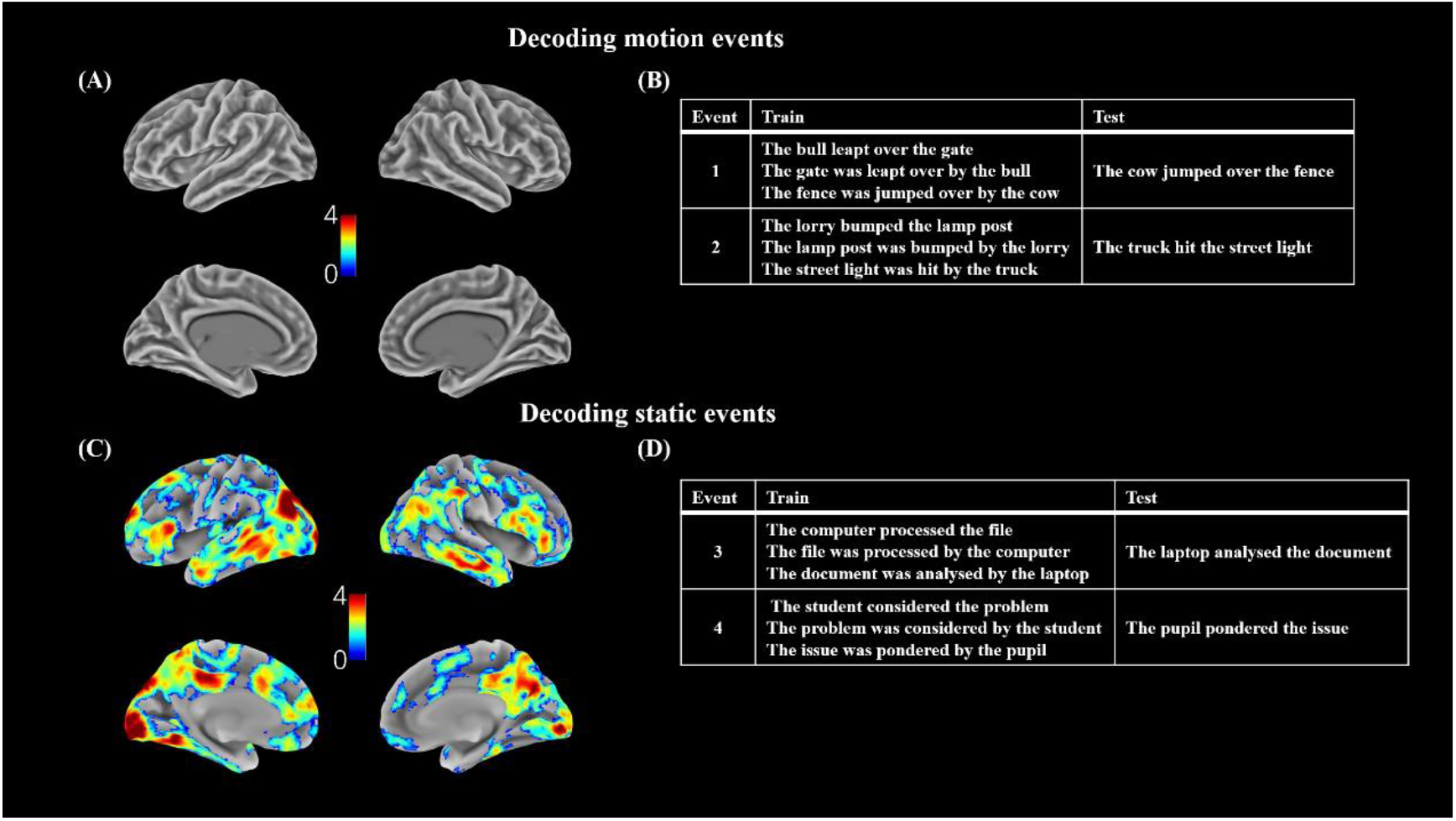
Decoding of specific motion and static events. (A) and (C) Decoding accuracy map, showing regions where classifier performance exceeded chance (corrected p < 0.05). (B) and (D) Examples of train and test patterns for one iteration of each analysis.

Figure 7 shows accuracies for each classifier for the LOTC regions of interest. When classifying the event type (motion/static), LOTC2 and LOTC3 showed significantly above-chance decoding accuracy, with highest accuracy in LOTC2. In contrast, none of the LOTC regions could successfully discriminate between the two different motion events. All regions could, however, distinguish between the two static events, in common with large swathes of temporal and occipital cortices (see Figure 6C).

**Figure 7.**
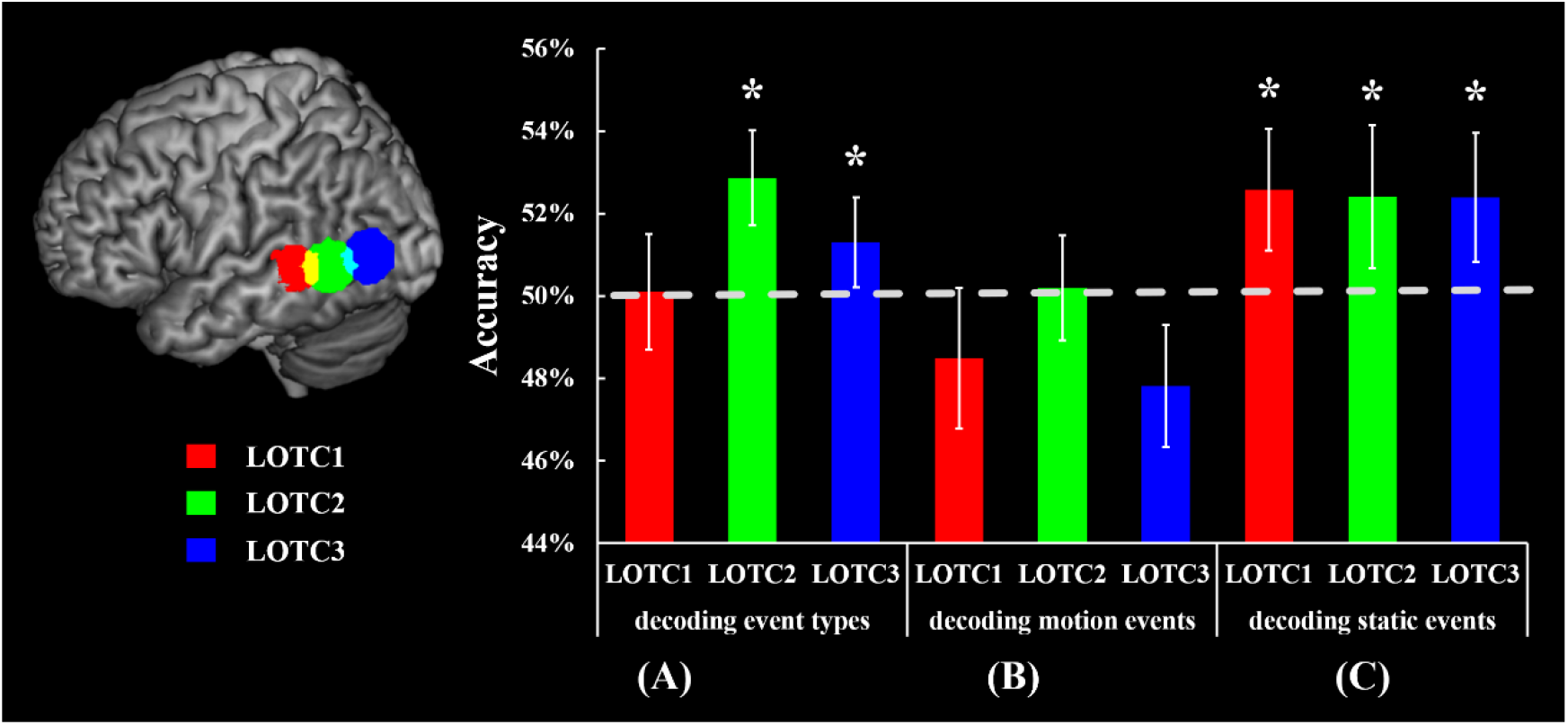
Decoding accuracies of LOTC1, LOTC2 and LOTC3 in different decoding models. Bars show one standard error of the mean.

Finally, as a control analysis, we took the searchlight classifier trained to discriminate the two motion events and tested its ability to categorize the static events. Since motion and static events do not correspond to one another, this attempt at classification should not be successful. As expected, no voxels showed above-chance decoding at our cluster-corrected threshold.

### PPI analyses

To investigate the functional connectivity of LOTC with other parts of the brain, PPI analyses were conducted using LOTC1, LOTC2 and LOTC3 as seed regions. Analyses tested for change in connectivity as a function of event type (motion vs. static). No regions showed significant increases in connectivity with LOTC for static relative to motion sentences. For motion events minus static events, however, significant effects were observed for the LOTC2 and LOTC3 seeds, as shown in Figure 8.

**Figure 8.**
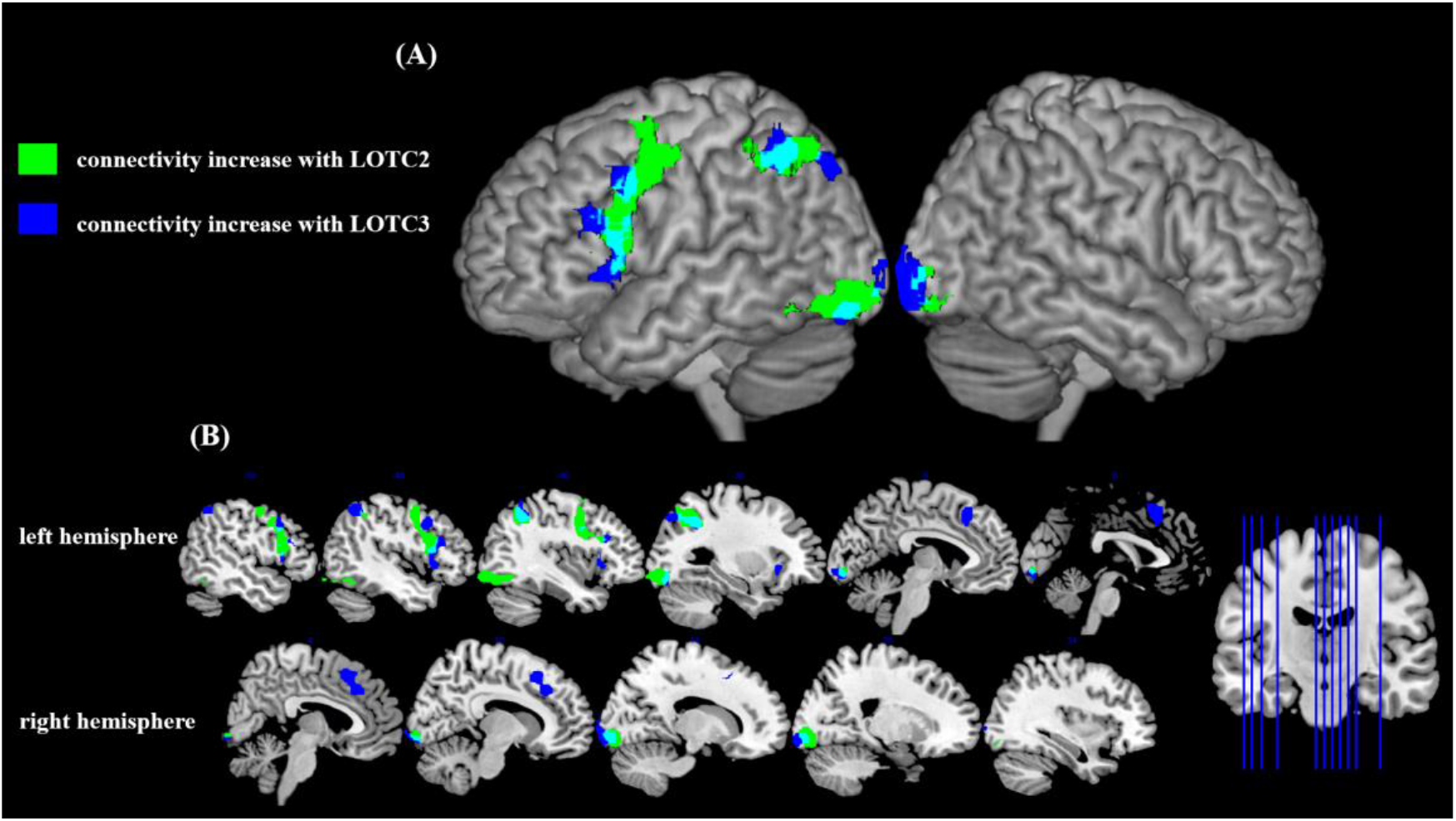
Regions showing increased connectivity with LOTC seeds for motion sentences vs. static sentences. (A) Surface render (cluster corrected p<0.05); (B) Slices (cluster corrected p<0.05)

A similar set of regions was revealed for both LOTC2 and LOTC3, including left precentral gyrus and lateral prefrontal cortex, left intraparietal sulcus, the presupplementary motor area and the lateral occipital cortex bilaterally. These areas all form part of a “multiple demand” network that shows increased engagement in response to cognitive challenges across multiple domains (Fedorenko et al., 2013). No regions showed significant connectivity changes for the LOTC1 seed.

## Discussion

LOTC has been implicated in conceptual processing of events involving motion but the precise nature and location of these semantic motion effects remains unclear. In this study, we used multiple neuroimaging analyses to examine LOTC’s role in motion concept representation. Participants made semantic decisions to sentences describing events that did or did not involve physical motion. MVPA revealed that activation patterns in LOTC discriminated between motion and static events. Significant decoding was observed in the middle and posterior parts of LOTC, although only the middle portion of the region showed an activation increase for motion sentences in univariate analyses. Moreover, PPI analyses indicated that the more posterior parts of LOTC increased their functional connectivity with the multiple demand network when participants processed motion sentences. Taken together, these results suggest that the more posterior parts of LOTC, very close to motion perception cortex, are most selectively involved in comprehending events involving motion, while the anterior part contributes to semantic processing in a more general fashion.

MVPA searchlight analyses across the whole brain indicated that activation patterns in left LOTC were most able to discriminate descriptions of static events from those that describe motion. This is consistent with the general view that this area of the cortex plays a particular role in comprehension, as well as direct perception, of motion events. Within LOTC, above-chance discrimination was observed in its middle and posterior areas, LOTC2 and LOTC3, with better decoding accuracy in LOTC2. In contrast, univariate contrasts of motion > static only found a significant effect in the middle part of LOTC. This difference may reflect the greater sensitivity of MVPA methods to subtle distinctions between conditions, which are not present in mean activation magnitude (Weaverdyck et al., 2020). Generally, the significant MVPA effects in LOTC suggest that its middle and posterior regions are sensitive to differences between motion and static events in language, including the posterior part that overlaps with motion perception cortex (V5).

PPI analyses also support the idea that posterior parts of LOTC are engaged when people process verbal descriptions of motion events. We used PPI to examine whether the functional connectivity of LOTC with other regions changed as a function of conceptual motion content. When processing motion events, the posterior LOTC2 and LOTC3 areas increased their connectivity with precentral gyrus, supplementary motor area (SMA), IPL and lateral occipital cortex. These regions form the multiple demand (MD) network which participates in domain-general cognitive control (De Baene et al., 2012; Koechlin et al., 2003; Kouneiher et al., 2009). This network has been implicated in controlled processing in domains such as semantic cognition (Jackson, 2021; Whitney et al., 2012), decision making (Coutlee & Huettel, 2012; Vickery & Jiang, 2009) and action coordination (Ridderinkhof et al., 2004; Rizzolatti et al., 2006). Here, we found that when participants made semantic decisions about motion events, these cognitive control areas showed increased interaction with the more posterior parts of LOTC. This could indicate that motion information encoded in LOTC was recruited as part of decision-making processes involved in the semantic judgements. In addition, the stronger connection between lateral visual cortex and LOTC might be caused not only by general cognitive control, but also by the mental imagery of motions. Compared to static events describing abstract actions (process, think), the motion events included verbs (jump over, hit) which were more likely to trigger imagination of dynamic images, thereby requiring more connectivity between lateral visual cortex and areas involved in perceiving motion.

The above results provide a more comprehensive view of LOTC involvement in processing motion concepts, on which previous studies hold different theories. Some researchers believe that motion concept representation requires re-enactment of perceptual experiences and the V5 would directly engage in this process (Hauk et al., 2004; Kiefer et al., 2012; Pulvermüller, 2005; Saygin et al., 2010). Others hold a weaker embodiment view, arguing that regions directly involved in perception do not necessarily engage in semantic processing, but areas close to them are recruited (Barsalou, 2003; Bedny et al., 2008; Kable et al., 2002; Martin & Chao, 2001). For understanding motion concepts, studies using univariate contrasts have found mixed evidence, with some reporting V5 involvement (Assmus et al., 2007; Glenberg & Kaschak, 2002; Revill et al., 2008; Rueschemeyer et al., 2010; Saygin et al., 2010) while others do not (Bedny et al., 2008; Gennari et al., 2007; Noppeney et al., 2005). In our study, we did not find any effect in LOTC3 in the univariate analyses but the more sensitive MVPA analyses did reveal motion effects, suggesting that motion perception regions are functionally involved in semantic processing. The LOTC2, as a region anterior to V5, also showed motion sensitivity in a range of analyses (univariate ROI analyses, MVPA and PPI), consistent with the weak embodiment view that areas anterior to V5 are sensitive to conceptual motion. Compared with middle and posterior parts of LOTC, the most anterior part, LOTC1, was not selective in its profile (no univariate motion effects, no PPI effects and no above-chance MVPA decoding) and we therefore conclude that the most anterior parts of LOTC play a more general role in semantic cognition. Overall, then, our results support general embodiment accounts of semantic processing, which implicate perceptual cortex and neighboring regions in processing related concepts. More importantly, the PPI analyses show how the various parts of LOTC coordinate with domain-general brain networks to process motion concepts. This complex interaction can be reflected more clearly in a study using a variety of analytic techniques, thus in future explorations, it is important to apply different analyses to develop a full picture.

MVPA models discriminating between the two motion events and the two static events revealed some unexpected results. The decoding model for the static events showed the expected pattern, with above-chance decoding in semantic regions such as prefrontal cortex, IFG and the anterior and posterior temporal lobes. However, when trained to discriminate between the two motion events, no regions showed above-chance performance and no effects were observed in LOTC (in searchlight or ROI analyses). Thus, although neural signals across the brain and in LOTC distinguished between motion and static sentences, they could not reliably discriminate between two different types of motion events.

For the static events decoding model, the generally high classification across many brain regions might relate to the subjects of the events, rather than the actions involved. The temoporoparetial junction region, lateral temporal cortex and posterior cingulate gyrus showed strong effects in discriminating between static events, and these areas are all parts of default mode network (DMN), which is engaged in processing socially relevant information and plays a key role for understanding others’ mental states (Li et al., 2014; Mars et al., 2012). One static event involved comprehending a person’s mental state and the other did not (computer processed file/student considered question), and DMN regions are likely to have responded differently to the human-relevant and object-relevant events for this reason.

The poor performance of LOTC in motion events decoding model was not predicted by embodied cognition theories, which would expect different patterns when different types of motion are described. One possible reason for this null result is the degree of perceptual simulation required by the task. Previous MVPA studies that have successfully decoded specific types of motion using neural responses in lateral posterior temporal cortex have elicited responses using videos of actions (Wurm et al., 2016; Wurm & Caramazza, 2019) or have asked people to make explicit action judgements about sentences (Wurm & Caramazza, 2019). In contrast, in the present study we used the relatively shallow task of asking participants to judge whether a written sentence was meaningful. The motion types might be decoded better if participants were required to deeply visualize different motions. However, this conclusion is speculative and further studies with more and deeper target stimuli are needed to investigate effects of different motion types in LOTC.

This study has a few limitations. Our two-category classification design required us to use a small number of events with simple sentence structure (agent-verb-patient). Future studies could use more events with different structures in their training sets, testing the degree to which our results generalize across conceptual and linguistic space. This is a particularly important point considering the limited ability of activation patterns to discriminate between the two motion events included in the present study.. Another possible limitation was that we did not know how deeply participants processed the sentences. The task goal may change people’s comprehension strategies, modulating the degree of embodiment and accompanying brain activity (for discussion, see Binder & Desai, 2011; Barsalou et al. 2008). For example, when people read novels, a detailed description of the environment and events could encourage people to enact detailed simulations of the content. In contrast, deciding whether a sentence is meaningful (like our experimental task) might be accomplished with less resort to mental imagery and simulation. Further research could ask participants to imagine the scene described by the sentences, to explore whether the LOTC’s function in motion concept representation would be affected by this factor.

In conclusion, using a range of analyses, this study found that middle and posterior parts of LOTC responded differently to motion and static events, including regions associated with perceptual processing of motion. We suggest that, future explorations combining activation-based, connectivity and pattern analysis techniques will be valuable in gaining further understanding of LOTC’s role in motion concept representation.

## Acknowledgements

PH was supported by a BBSRC grant (BB/T004444/1). Imaging was carried out at the Edinburgh Imaging Facility (www.ed.ac.uk/edinburgh-imaging), University of Edinburgh, which is part of the SINAPSE collaboration (www.sinapse.ac.uk). We are grateful to the University of Minnesota Center for Magnetic Resonance Research for sharing their neuroimaging sequences.

## Reference

Ashburner, J. (2007). A fast diffeomorphic image registration algorithm. Neuroimage, 38(1), 95–113. https://doi.org/DOI10.1016/j.neuroimage.2007.07.007

Ashburner, J., Barnes, G., Chen, C.-C., Daunizeau, J., Flandin, G., Friston, K., Kiebel, S., Kilner, J., Litvak, V., & Moran, R. (2014). SPM12 manual. Wellcome Trust Centre for Neuroimaging, London, UK, 2464.

Assmus, A., Giessing, C., Weiss, P. H., & Fink, G. R. (2007). Functional interactions during the retrieval of conceptual action knowledge: an fMRI study. Journal of cognitive neuroscience, 19(6), 1004–1012.

Asyraff, A., Lemarchand, R., Tamm, A., & Hoffman, P. (2021). Stimulus-independent neural coding of event semantics: Evidence from cross-sentence fMRI decoding. Neuroimage, 236, 118073.

Barron, H. C., Garvert, M. M., & Behrens, T. E. (2016). Repetition suppression: a means to index neural representations using BOLD? Philosophical Transactions of the Royal Society B: Biological Sciences, 371(1705), 20150355.

Barsalou, L. (2003). Situated simulation in the human conceptual system. Language and cognitive processes, 18(5-6), 513–562.

Barsalou, L. W. (1999). Perceptual symbol systems. Behavioral and brain sciences, 22(4), 577–660.

Beauchamp, M. S., Lee, K. E., Argall, B. D., & Martin, A. (2004). Integration of auditory and visual information about objects in superior temporal sulcus. Neuron, 41(5), 809–823.

Beauchamp, M. S., Lee, K. E., Haxby, J. V., & Martin, A. (2002). Parallel visual motion processing streams for manipulable objects and human movements. Neuron, 34(1), 149–159.

Bedny, M., & Caramazza, A. (2011). Perception, action, and word meanings in the human brain: the case from action verbs. Annals of the New York Academy of Sciences, 1224(1), 81–95.

Bedny, M., Caramazza, A., Grossman, E., Pascual-Leone, A., & Saxe, R. (2008). Concepts are more than percepts: the case of action verbs. Journal of Neuroscience, 28(44), 11347–11353.

Bedny, M., Caramazza, A., Pascual-Leone, A., & Saxe, R. (2012). Typical neural representations of action verbs develop without vision. Cerebral cortex, 22(2), 286–293.

Bedny, M., Dravida, S., & Saxe, R. (2014). Shindigs, brunches, and rodeos: The neural basis of event words. Cognitive, Affective, & Behavioral Neuroscience, 14(3), 891–901.

Beilock, S. L., Lyons, I. M., Mattarella-Micke, A., Nusbaum, H. C., & Small, S. L. (2008). Sports experience changes the neural processing of action language. Proceedings of the National Academy of Sciences, 105(36), 13269–13273.

Brambati, S. M., Myers, D., Wilson, A., Rankin, K. P., Allison, S. C., Rosen, H. J., Miller, B. L., & Gorno-Tempini, M. L. (2006). The anatomy of category-specific object naming in neurodegenerative diseases. Journal of cognitive neuroscience, 18(10), 1644–1653.

Brysbaert, M., Warriner, A. B., & Kuperman, V. (2014). Concreteness ratings for 40 thousand generally known English word lemmas. Behavior research methods, 46(3), 904–911.

Bucur, M., & Papagno, C. (2021). An ALE meta-analytical review of the neural correlates of abstract and concrete words. Scientific reports, 11(1), 1–24.

Buxbaum, L. J., Shapiro, A. D., & Coslett, H. B. (2014). Critical brain regions for tool-related and imitative actions: a componential analysis. Brain, 137(7), 1971–1985.

Calvert, G. A., Brammer, M. J., Bullmore, E. T., Campbell, R., Iversen, S. D., & David, A. S. (1999). Response amplification in sensory-specific cortices during crossmodal binding. NeuroReport, 10(12), 2619–2623.

Calvert, G. A., Campbell, R., & Brammer, M. J. (2000). Evidence from functional magnetic resonance imaging of crossmodal binding in the human heteromodal cortex. Current Biology, 10(11), 649–657.

Chen, E., Widick, P., & Chatterjee, A. (2008). Functional–anatomical organization of predicate metaphor processing. Brain and language, 107(3), 194–202.

Coutlee, C. G., & Huettel, S. A. (2012). The functional neuroanatomy of decision making: prefrontal control of thought and action. Brain Research, 1428, 3–12.

Cross, E. S., Hamilton, A. F. d. C., & Grafton, S. T. (2006). Building a motor simulation de novo: observation of dance by dancers. Neuroimage, 31(3), 1257–1267.

De Baene, W., Albers, A. M., & Brass, M. (2012). The what and how components of cognitive control. Neuroimage, 63(1), 203–211.

Decety, L., & Grezes, J. (2006). Multiple perspectives on the psychological and neural bases of understanding other people’s behavior. Brain Research, 1079, 4–14.

Desai, R. H., Conant, L. L., Binder, J. R., Park, H., & Seidenberg, M. S. (2013). A piece of the action: modulation of sensory-motor regions by action idioms and metaphors. Neuroimage, 83, 862–869.

Dravida, S., Saxe, R., & Bedny, M. (2013). People can understand descriptions of motion without activating visual motion brain regions. Front Psychol, 4, 537. https://doi.org/10.3389/fpsyg.2013.00537

Dumoulin, S. O., Bittar, R. G., Kabani, N. J., Baker Jr, C. L., Le Goualher, G., Pike, G. B., & Evans, A. C. (2000). A new anatomical landmark for reliable identification of human area V5/MT: a quantitative analysis of sulcal patterning. Cerebral cortex, 10(5), 454–463.

DuPre, E., Salo, T., Markello, R., Kundu, P., Whitaker, K., & Handwerker, D. (2019). ME-ICA/tedana: 0.0. 6. Zenodo.

Fedorenko, E., Duncan, J., & Kanwisher, N. (2013). Broad domain generality in focal regions of frontal and parietal cortex. Proceedings of the National Academy of Sciences, 110(41), 16616–16621.

Feinberg, D. A., Moeller, S., Smith, S. M., Auerbach, E., Ramanna, S., Glasser, M. F., Miller, K. L., Ugurbil, K., & Yacoub, E. (2010). Multiplexed echo planar imaging for sub-second whole brain FMRI and fast diffusion imaging. PloS one, 5(12), e15710.

Friston, K., Buechel, C., Fink, G., Morris, J., Rolls, E., & Dolan, R. J. (1997). Psychophysiological and modulatory interactions in neuroimaging. Neuroimage, 6(3), 218–229.

Gallese, V., & Lakoff, G. (2005). The brain’s concepts: The role of the sensory-motor system in conceptual knowledge. Cognitive neuropsychology, 22(3-4), 455–479.

Gardumi, A., Ivanov, D., Hausfeld, L., Valente, G., Formisano, E., & Uludağ, K. (2016). The effect of spatial resolution on decoding accuracy in fMRI multivariate pattern analysis. Neuroimage, 132, 32–42.

Gennari, S. P. (2012). Representing motion in language comprehension: lessons from neuroimaging. Language and Linguistics Compass, 6(2), 67–84.

Gennari, S. P., MacDonald, M. C., Postle, B. R., & Seidenberg, M. S. (2007). Context-dependent interpretation of words: Evidence for interactive neural processes. Neuroimage, 35(3), 1278–1286.

Gitelman, D. R., Penny, W. D., Ashburner, J., & Friston, K. J. (2003). Modeling regional and psychophysiologic interactions in fMRI: the importance of hemodynamic deconvolution. Neuroimage, 19(1), 200–207.

Glenberg, A. M., & Kaschak, M. P. (2002). Grounding language in action. Psychonomic bulletin & review, 9(3), 558–565.

Hauk, O., Johnsrude, I., & Pulvermüller, F. (2004). Somatotopic representation of action words in human motor and premotor cortex. Neuron, 41(2), 301–307.

Hendriks, M. H., Daniels, N., Pegado, F., & Op de Beeck, H. P. (2017). The effect of spatial smoothing on representational similarity in a simple motor paradigm. Frontiers in neurology, 8, 222.

Hoeren, M., Kümmerer, D., Bormann, T., Beume, L., Ludwig, V. M., Vry, M.-S., Mader, I., Rijntjes, M., Kaller, C. P., & Weiller, C. (2014). Neural bases of imitation and pantomime in acute stroke patients: distinct streams for praxis. Brain, 137(10), 2796–2810.

Humphreys, G. F., Newling, K., Jennings, C., & Gennari, S. P. (2013). Motion and actions in language: semantic representations in occipito-temporal cortex. Brain Lang, 125(1), 94–105. https://doi.org/10.1016/j.bandl.2013.01.008

Jackson, R. L. (2021). The neural correlates of semantic control revisited. Neuroimage, 224, 117444.

Kable, J. W., Kan, I. P., Wilson, A., Thompson-Schill, S. L., & Chatterjee, A. (2005). Conceptual representations of action in the lateral temporal cortex. Journal of cognitive neuroscience, 17(12), 1855–1870.

Kable, J. W., Lease-Spellmeyer, J., & Chatterjee, A. (2002). Neural substrates of action event knowledge. Journal of cognitive neuroscience, 14(5), 795–805.

Kalénine, S., Buxbaum, L. J., & Coslett, H. B. (2010). Critical brain regions for action recognition: lesion symptom mapping in left hemisphere stroke. Brain, 133(11), 3269–3280.

Kemmerer, D., Rudrauf, D., Manzel, K., & Tranel, D. (2012). Behavioral patterns and lesion sites associated with impaired processing of lexical and conceptual knowledge of actions. Cortex, 48(7), 826–848.

Kiefer, M., Trumpp, N., Herrnberger, B., Sim, E.-J., Hoenig, K., & Pulvermüller, F. (2012). Dissociating the representation of action-and sound-related concepts in middle temporal cortex. Brain and language, 122(2), 120–125.

Koechlin, E., Ody, C., & Kouneiher, F. (2003). The architecture of cognitive control in the human prefrontal cortex. Science, 302(5648), 1181–1185.

Kouneiher, F., Charron, S., & Koechlin, E. (2009). Motivation and cognitive control in the human prefrontal cortex. Nature neuroscience, 12(7), 939–945.

Kourtzi, Z., & Kanwisher, N. (2000). Activation in human MT/MST by static images with implied motion. Journal of cognitive neuroscience, 12(1), 48–55.

Kundu, P., Voon, V., Balchandani, P., Lombardo, M. V., Poser, B. A., & Bandettini, P. A. (2017). Multi-echo fMRI: a review of applications in fMRI denoising and analysis of BOLD signals. Neuroimage, 154, 59–80.

Li, W., Mai, X., & Liu, C. (2014). The default mode network and social understanding of others: what do brain connectivity studies tell us. Frontiers in human neuroscience, 74.

Lingnau, A., & Downing, P. E. (2015). The lateral occipitotemporal cortex in action. Trends Cogn Sci, 19(5), 268–277. https://doi.org/10.1016/j.tics.2015.03.006

Lingnau, A., & Petris, S. (2013). Action understanding within and outside the motor system: the role of task difficulty. Cerebral cortex, 23(6), 1342–1350.

Liu, L. D., Haefner, R. M., & Pack, C. C. (2016). A neural basis for the spatial suppression of visual motion perception. Elife, 5, e16167.

Malikovic, A., Amunts, K., Schleicher, A., Mohlberg, H., Eickhoff, S. B., Wilms, M., Palomero-Gallagher, N., Armstrong, E., & Zilles, K. (2007). Cytoarchitectonic analysis of the human extrastriate cortex in the region of V5/MT+: a probabilistic, stereotaxic map of area hOc5. Cerebral cortex, 17(3), 562–574.

Mars, R. B., Neubert, F.-X., Noonan, M. P., Sallet, J., Toni, I., & Rushworth, M. F. (2012). On the relationship between the “default mode network” and the “social brain”. Frontiers in human neuroscience, 6, 189.

Martin, A., & Chao, L. L. (2001). Semantic memory and the brain: structure and processes. Current opinion in neurobiology, 11(2), 194–201.

Martin, A., Haxby, J. V., Lalonde, F. M., Wiggs, C. L., & Ungerleider, L. G. (1995). Discrete cortical regions associated with knowledge of color and knowledge of action. Science, 270(5233), 102–105.

Moeller, S., Yacoub, E., Olman, C. A., Auerbach, E., Strupp, J., Harel, N., & Uğurbil, K. (2010). Multiband multislice GE-EPI at 7 tesla, with 16-fold acceleration using partial parallel imaging with application to high spatial and temporal whole-brain fMRI. Magnetic resonance in medicine, 63(5), 1144–1153.

Mumford, J. A., Davis, T., & Poldrack, R. A. (2014). The impact of study design on pattern estimation for single-trial multivariate pattern analysis. Neuroimage, 103, 130–138.

Noppeney, U., Josephs, O., Kiebel, S., Friston, K. J., & Price, C. J. (2005). Action selectivity in parietal and temporal cortex. Cognitive Brain Research, 25(3), 641–649.

O’Reilly, J. X., Woolrich, M. W., Behrens, T. E., Smith, S. M., & Johansen-Berg, H. (2012). Tools of the trade: psychophysiological interactions and functional connectivity. Social cognitive and affective neuroscience, 7(5), 604–609.

Oosterhof, N. N., Connolly, A. C., & Haxby, J. V. (2016). CoSMoMVPA: multi-modal multivariate pattern analysis of neuroimaging data in Matlab/GNU Octave. Frontiers in neuroinformatics, 10, 27.

Peelen, M. V., Romagno, D., & Caramazza, A. (2012). Independent representations of verbs and actions in left lateral temporal cortex. Journal of cognitive neuroscience, 24(10), 2096–2107.

Power, J. D., Plitt, M., Kundu, P., Bandettini, P. A., & Martin, A. (2017). Temporal interpolation alters motion in fMRI scans: Magnitudes and consequences for artifact detection. PloS one, 12(9), e0182939.

Pulvermüller, F. (2005). Brain mechanisms linking language and action. Nature reviews neuroscience, 6(7), 576–582.

Revill, K. P., Aslin, R. N., Tanenhaus, M. K., & Bavelier, D. (2008). Neural correlates of partial lexical activation. Proceedings of the National Academy of Sciences, 105(35), 13111–13115.

Ridderinkhof, K. R., Van Den Wildenberg, W. P., Segalowitz, S. J., & Carter, C. S. (2004). Neurocognitive mechanisms of cognitive control: the role of prefrontal cortex in action selection, response inhibition, performance monitoring, and reward-based learning. Brain and cognition, 56(2), 129–140.

Rizzolatti, G., Ferrari, P. F., Rozzi, S., & Fogassi, L. (2006). The inferior parietal lobule: where action becomes perception. Novartis Foundation Symposium

Rueschemeyer, S.-A., Glenberg, A. M., Kaschak, M., Mueller, K., & Friederici, A. (2010). Top-down and bottom-up contributions to understanding sentences describing objects in motion. Frontiers in Psychology, 1, 183.

Saygin, A. P., McCullough, S., Alac, M., & Emmorey, K. (2010). Modulation of BOLD response in motion-sensitive lateral temporal cortex by real and fictive motion sentences. Journal of cognitive neuroscience, 22(11), 2480–2490.

Schultz, J., Friston, K. J., O’Doherty, J., Wolpert, D. M., & Frith, C. D. (2005). Activation in posterior superior temporal sulcus parallels parameter inducing the percept of animacy. Neuron, 45(4), 625–635.

Senior, C., Barnes, J., Giampietroc, V., Simmons, A., Bullmore, E., Brammer, M., & David, A. (2000). The functional neuroanatomy of implicit-motion perception or ‘representational momentum’. Current Biology, 10(1), 16–22.

Smith, S. M., & Nichols, T. E. (2009). Threshold-free cluster enhancement: addressing problems of smoothing, threshold dependence and localisation in cluster inference. Neuroimage, 44(1), 83–98.

Stelzer, J., Chen, Y., & Turner, R. (2013). Statistical inference and multiple testing correction in classification-based multi-voxel pattern analysis (MVPA): random permutations and cluster size control. Neuroimage, 65, 69–82.

Thompson, J. C., Clarke, M., Stewart, T., & Puce, A. (2005). Configural processing of biological motion in human superior temporal sulcus. Journal of Neuroscience, 25(39), 9059–9066.

Todd, M. T., Nystrom, L. E., & Cohen, J. D. (2013). Confounds in multivariate pattern analysis: theory and rule representation case study. Neuroimage, 77, 157–165.

Tootell, R. B., Reppas, J. B., Kwong, K. K., Malach, R., Born, R. T., Brady, T. J., Rosen, B. R., & Belliveau, J. W. (1995). Functional analysis of human MT and related visual cortical areas using magnetic resonance imaging. Journal of Neuroscience, 15(4), 3215–3230.

Tucciarelli, R., Wurm, M., Baccolo, E., & Lingnau, A. (2019). The representational space of observed actions. Elife, 8. https://doi.org/10.7554/eLife.47686

Urgesi, C., Candidi, M., & Avenanti, A. (2014). Neuroanatomical substrates of action perception and understanding: an anatomic likelihood estimation meta-analysis of lesion-symptom mapping studies in brain injured patients. Frontiers in human neuroscience, 8, 344.

Van Heuven, W. J., Mandera, P., Keuleers, E., & Brysbaert, M. (2014). SUBTLEX-UK: A new and improved word frequency database for British English. Quarterly journal of experimental psychology, 67(6), 1176–1190.

Vickery, T. J., & Jiang, Y. V. (2009). Inferior parietal lobule supports decision making under uncertainty in humans. Cerebral cortex, 19(4), 916–925.

Wall, M. B., Lingnau, A., Ashida, H., & Smith, A. T. (2008). Selective visual responses to expansion and rotation in the human MT complex revealed by functional magnetic resonance imaging adaptation. European Journal of Neuroscience, 27(10), 2747–2757.

Wallentin, M., Lund, T. E., Östergaard, S., Östergaard, L., & Roepstorff, A. (2005). Motion verb sentences activate left posterior middle temporal cortex despite static context. NeuroReport, 16(6), 649–652.

Wang, J., Conder, J. A., Blitzer, D. N., & Shinkareva, S. V. (2010). Neural representation of abstract and concrete concepts: A meta-analysis of neuroimaging studies. Human brain mapping, 31(10), 1459–1468.

Watson, C. E., Cardillo, E. R., Ianni, G. R., & Chatterjee, A. (2013). Action concepts in the brain: an activation likelihood estimation meta-analysis. J Cogn Neurosci, 25(8), 1191–1205. https://doi.org/10.1162/jocn_a_00401

Weaverdyck, M. E., Lieberman, M. D., & Parkinson, C. (2020). Tools of the Trade Multivoxel pattern analysis in fMRI: a practical introduction for social and affective neuroscientists. Soc Cogn Affect Neurosci, 15(4), 487–509.

Weiner, K. S., & Grill-Spector, K. (2013). Neural representations of faces and limbs neighbor in human high-level visual cortex: evidence for a new organization principle. Psychological research, 77(1), 74–97.

Whitney, C., Kirk, M., O’Sullivan, J., Lambon Ralph, M. A., & Jefferies, E. (2012). Executive semantic processing is underpinned by a large-scale neural network: revealing the contribution of left prefrontal, posterior temporal, and parietal cortex to controlled retrieval and selection using TMS. Journal of cognitive neuroscience, 24(1), 133–147.

Woolgar, A., Golland, P., & Bode, S. (2014). Coping with confounds in multivoxel pattern analysis: What should we do about reaction time differences? A comment on Todd, Nystrom & Cohen 2013. Neuroimage, 98, 506–512.

Wurm, M. F., Ariani, G., Greenlee, M. W., & Lingnau, A. (2016). Decoding Concrete and Abstract Action Representations During Explicit and Implicit Conceptual Processing. Cereb Cortex, 26(8), 3390–3401. https://doi.org/10.1093/cercor/bhv169

Wurm, M. F., & Caramazza, A. (2019). Distinct roles of temporal and frontoparietal cortex in representing actions across vision and language. Nat Commun, 10(1), 289. https://doi.org/10.1038/s41467-018-08084-y

Wurm, M. F., Caramazza, A., & Lingnau, A. (2017). Action Categories in Lateral Occipitotemporal Cortex Are Organized Along Sociality and Transitivity. J Neurosci, 37(3), 562–575. https://doi.org/10.1523/JNEUROSCI.1717-16.2016

Xu, J., Moeller, S., Auerbach, E. J., Strupp, J., Smith, S. M., Feinberg, D. A., Yacoub, E., & Uğurbil, K. (2013). Evaluation of slice accelerations using multiband echo planar imaging at 3 T. Neuroimage, 83, 991–1001.

Yarkoni, T., Poldrack, R. A., Nichols, T. E., Van Essen, D. C., & Wager, T. D. (2011). Large-scale automated synthesis of human functional neuroimaging data. Nature methods, 8(8), 665–670.

Zeki, S., Watson, J., Lueck, C., Friston, K. J., Kennard, C., & Frackowiak, R. (1991). A direct demonstration of functional specialization in human visual cortex. Journal of Neuroscience, 11(3), 641–649.

