## Supplementary for "Representation of motion concepts in occipitotemporal cortex: fMRI activation, decoding and connectivity analyses"

Supplementary Table 1: List of Stimuli

| Sentence | Condition | Meaningfulness | Syntax Form | Lexical Form | Event |
| --- | --- | --- | --- | --- | --- |
| The bull leapt over the gate. | motion | meaningful | Active | 1 | Event1 |
| The gate was leapt over by the bull. | motion | meaningful | Passive | 1 | Event1 |
| The cow jumped over the fence. | motion | meaningful | Active | 2 | Event1 |
| The fence was jumped over by the cow. | motion | meaningful | Passive | 2 | Event1 |
| The lorry bumped the lamp post. | motion | meaningful | Active | 1 | Event2 |
| The lamp post was bumped by the lorry. | motion | meaningful | Passive | 1 | Event2 |
| The truck hit the street light. | motion | meaningful | Active | 2 | Event2 |
| The street light was hit by the truck. | motion | meaningful | Passive | 2 | Event2 |
| The computer processed the file. | static | meaningful | Active | 1 | Event3 |
| The file was processed by the computer. | static | meaningful | Passive | 1 | Event3 |
| The laptop analysed the document. | static | meaningful | Active | 2 | Event3 |
| The document was analysed by the laptop. | static | meaningful | Passive | 2 | Event3 |
| The student considered the problem. | static | meaningful | Active | 1 | Event4 |
| The problem was considered by the student. | static | meaningful | Passive | 1 | Event4 |
| The pupil pondered the issue. | static | meaningful | Active | 2 | Event4 |
| The issue was pondered by the pupil. | static | meaningful | Passive | 2 | Event4 |
| The files pondered the truck | control | anomalous | Active | 1+2 |  |
| The streetlight was jumped over by the computer | control | anomalous | Passive | 1+2 |  |
| The problem hit the cow | control | anomalous | Active | 1+2 |  |
| The document was considered by the lorry | control | anomalous | Passive | 1+2 |  |
| The gate pondered the pupil | control | anomalous | Active | 1+2 |  |
| The problem was hit by the cow | control | anomalous | Passive | 1+2 |  |
| The computer jumped over the streetlight | control | anomalous | Active | 1+2 |  |
| The fence was analysed by the cow | control | anomalous | Passive | 1+2 |  |
| The issue bumped the bull | control | anomalous | Active | 1+2 |  |
| The issue was bumped by the bull | control | anomalous | Passive | 1+2 |  |
| The laptop leapt over the streetlight | control | anomalous | Active | 1+2 |  |
| The gate was pondered by the pupil | control | anomalous | Passive | 1+2 |  |
| The fence processed the student | control | anomalous | Active | 1+2 |  |
| The truck was pondered by the files | control | anomalous | Passive | 1+2 |  |
| The streetlight was leapt over by the laptop | control | anomalous | Passive | 1+2 |  |
| The lorry was considered by the document | control | anomalous | Passive | 1+2 |  |

Supplementary Table 2: Mean properties for each event

| Event | Condition | Concreteness | Frequency | Vision | Motion | Sound | Emotion |
| --- | --- | --- | --- | --- | --- | --- | --- |
| Event1 | Motion | 3.02 | 5.875 | 5.359 | 5.532 | 3.566 | 2.207 |
| Event2 | Motion | 3.068 | 5.508 | 5.499 | 5.312 | 4.514 | 2.953 |
| Event3 | Static | 2.775 | 5.553 | 2.913 | 2.162 | 2.023 | 1.619 |
| Event4 | Static | 2.303 | 5.88 | 3.489 | 2.313 | 2.228 | 3.065 |
| Mean Difference  (Motion-Static) | | 0.505 | -0.025 | 2.228 | 3.184 | 1.915 | 0.24 |
| Motion Mean | | 3.04 | 5.69 | 5.43 | 5.42 | 4.04 | 2.58 |
| Static Mean | | 2.54 | 5.72 | 3.2 | 2.24 | 2.13 | 2.34 |
| T-value(Motion VS Static) | | 3.654** | -0.146 | 11.868*** | 17.763*** | 7.836*** | 0.67 |
| df | | 13.27 | 11.77 | 13.23 | 9.66 | 7.99 | 12.23 |

** = *p* < 0.01; *** = *p* < 0.001.

Supplementary Table 3: Mean properties for each target sentence

| Sentence | Condition | Conc | Freq | Vision | Motion | Sound | Emotion |
| --- | --- | --- | --- | --- | --- | --- | --- |
| The bull leapt over the gate | Motion | 3.17 | 5.7 | 5.65 | 5.87 | 3.87 | 2.57 |
| The gate was leapt over by the bull | Motion | 2.78 | 5.92 | 4.96 | 4.91 | 3.48 | 2.35 |
| The cow jumped over the fence | Motion | 3.27 | 5.85 | 5.78 | 5.91 | 3.65 | 2.09 |
| The fence was jumped over by the cow | Motion | 2.86 | 6.03 | 5.04 | 5.43 | 3.26 | 1.83 |
| The lorry bumped the lamp post | Motion | 3.29 | 5.27 | 5.43 | 5.09 | 4.35 | 2.61 |
| The lamp post was bumped by the lorry | Motion | 2.81 | 5.64 | 4.87 | 4.78 | 3.74 | 2.43 |
| The truck hit the streetlight | Motion | 3.33 | 5.39 | 5.95 | 5.95 | 5.14 | 3.55 |
| The streetlight was hit by the truck | Motion | 2.84 | 5.73 | 5.74 | 5.43 | 4.83 | 3.22 |
| The computer processed the file | Static | 3.03 | 5.82 | 2.87 | 2.09 | 2.22 | 1.61 |
| The file was processed by the computer | Static | 2.62 | 6.04 | 3.04 | 2.52 | 2.04 | 1.61 |
| The laptop analyzed the document | Static | 2.91 | 4.94 | 2.83 | 2.04 | 1.87 | 1.65 |
| The document was analyzed by the laptop | Static | 2.54 | 5.41 | 2.91 | 2 | 1.96 | 1.61 |
| The pupil pondered the issue | Static | 2.4 | 5.5 | 3.65 | 2.3 | 2.43 | 3.43 |
| The issue was pondered by the student | Static | 2.18 | 5.81 | 3.48 | 2.43 | 2.09 | 3.22 |
| The student considered the problem | Static | 2.43 | 6.03 | 3.52 | 2.43 | 2.26 | 3.3 |
| The problem was considered by the student | Static | 2.2 | 6.18 | 3.3 | 2.09 | 2.13 | 2.3 |

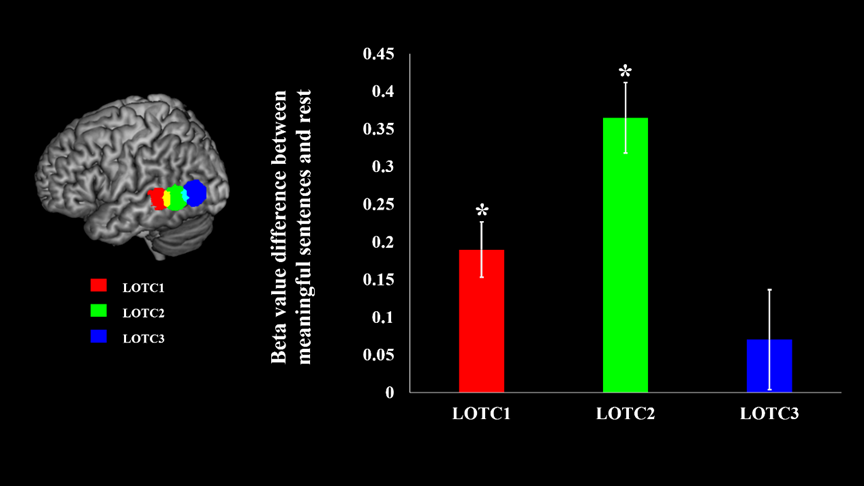

*Figure 1. The 3 ROIs’ beta value differences between ‘meaningful’ and ‘rest’*
